# Simulated low vision disrupts geometry-dependent contact-point selection during grasping

**DOI:** 10.64898/2026.11.24.769076

**Authors:** Carla Lanca, Zoltan Derzsi, Robert Volcic

**Affiliations:** Division of Science, New York University Abu Dhabi, Abu Dhabi, UAE; Center for Brain and Health, New York University Abu Dhabi, Abu Dhabi, UAE; Center for Artificial Intelligence and Robotics, New York University Abu Dhabi, Abu Dhabi, UAE

**Keywords:** simulated low vision, visually guided action, grasping, contact-point selection, object manipulation, visuomotor control

## Abstract

**Purpose:** To investigate how simulated low vision (SV) affects visually guided grasping, focusing on contact-point selection relative to object geometry.

**Methods:** Twenty-three adults performed reach-to-grasp movements toward 3D-printed objects under normal-vision (NV) and SV conditions. SV was induced using Cambridge Simulation Glasses with four diffusing filter layers. Participants grasped objects with alternating convex and concave regions presented at different orientations and two contrast levels while finger movements were recorded using motion tracking. Bayesian hierarchical analyses examined reach transport, grasp formation and its scaling, and grasp placement relative to object geometry.

**Results:** Compared with NV, movements under SV followed shorter trajectories (7.3 mm; 95% CrI, 1.4–13.0 mm) and lower finger heights (5.0 mm; 95% CrI, 1.3–8.7 mm). SV caused maximum grip aperture to occur earlier by 27 ms (95% CrI, 1.0–51.7 ms). Contact points were positioned farther from the object’s center of mass (0.64 mm; 95% CrI, 0.11–1.17 mm). Under NV, grasp axes clustered near the centers of convex and concave regions, whereas under SV, selections approached a random distribution, particularly at low contrast. Under SV, contact-point selection variability increased by 3.91° (95% CrI, 2.55–5.20°) for concave regions and 4.74° (95% CrI, 3.45–6.04°) for convex regions. However, the overall proportion of convex and concave grasp-point selections remained similar across visual conditions.

**Conclusions:** SV altered how visual information was used to guide interactions with object structure without disrupting successful grasp performance. These findings suggest that visual impairment may affect functional object manipulation in ways not captured by conventional measures of visual performance.

## 1 Introduction

Visual information guides hand movements in many everyday activities, including picking up objects and handling tools. During reach-to-grasp movements, visual information about object size, shape, orientation, and spatial layout is used to plan and perform hand movements, select appropriate contact points, and maintain grasp stability throughout the action.^1–4^ Although grasping appears effortless, successful performance relies on complex visuomotor processes that continuously integrate sensory information with motor commands. The way an object is grasped is determined by a combination of factors, including the physical properties of the object, task demands, and the quality of the available visual information.^5–12^

Visual information supports both initial planning and online control of grasping. During reach-to-grasp actions, visual and proprioceptive feedback are continuously integrated to guide digit placement and achieve stable interactions with the object. Unexpected changes in object properties during movement execution elicit rapid grasp adjustments, indicating that visuomotor control depends on dynamic visual processing rather than solely on pre-planned motor commands.^13–18^

The importance of visual input is further illustrated by studies of individuals with visual impairment. Reach-to-grasp performance is often impaired in populations with age-related macular degeneration,^19,20^ glaucoma,^21,22^ and amblyopia or reduced stereopsis.^23–25^ Degraded vision consistently causes slower movements, longer movement durations, increased reliance on online adjustments, and impaired movement planning, particularly under reduced object contrast.^26–28^ However, the effects on core kinematic parameters such as maximum grip aperture and movement velocity appear to vary across studies, likely depending on the nature and severity of visual degradation and task demands.^20,22,29,30^

Beyond these temporal and kinematic aspects, successful grasping behavior ultimately depends on the selection of stable contact points on an object’s surface. Previous work has shown that grasp-point selection reflects interactions between perceptual and biomechanical constraints, including preferred grasp orientation and grasp stability requirements.^31,32^ Although many potential contact points exist, only a limited subset yields effective grasps, highlighting the systematic nature of contact-point selection.^9^

Grasp stability requires integrating two spatial constraints: aligning the grasp axis with the object’s center of mass and anchoring the digits onto suitable local geometry.^5,33–41^ Despite the critical role of this spatial organization, how degraded vision affects contact-point selection remains largely unexplored. Achieving both spatial requirements simultaneously might become particularly challenging under degraded visual conditions, where extraction of fine structural cues is impaired.

While clinical impairments vary, experimentally induced blur provides a controlled approach to investigate how such visual deficits influence visuomotor behavior.^30,42–45^ By reducing visual acuity and causing contrast attenuation, blur limits the availability of critical visual cues regarding object boundaries and surface structure, potentially affecting the accuracy and precision of visually guided actions.

The current study investigated how degraded vision influences the spatial organization of a grasp. While we evaluated standard temporal and kinematic parameters, our specific focus was on how blur induced by diffusing filters and object contrast alter center-of-mass alignment and influences contact-point selection based on local object geometry. Understanding how visual degradation affects contact-point selection may therefore uncover functional difficulties in visually guided actions that conventional clinical measures do not fully capture.

## 2 Materials and Methods

### 2.1 Participants

Twenty-three right-handed participants took part in this study (mean ± SD age: 21 ± 2.4 years). All had normal or corrected-to-normal vision and no history of neurological disorders. Participants were näıve to the study purpose and were provided with a subsistence allowance. Participants provided informed written consent and the study was approved by the New York University Abu Dhabi Institutional Review Board.

### 2.2 Apparatus and procedure

Participants sat at a table with their right thumb and index positioned on a start location aligned with the body midline. Following an auditory cue, participants reached toward the target object, grasped it using a precision grip, lifted it without removing it from the holder, returned it to the holder, and moved the hand back to the start position (Figure 1A). Movements were performed at a natural speed without time constraints.

**Fig. 1.**
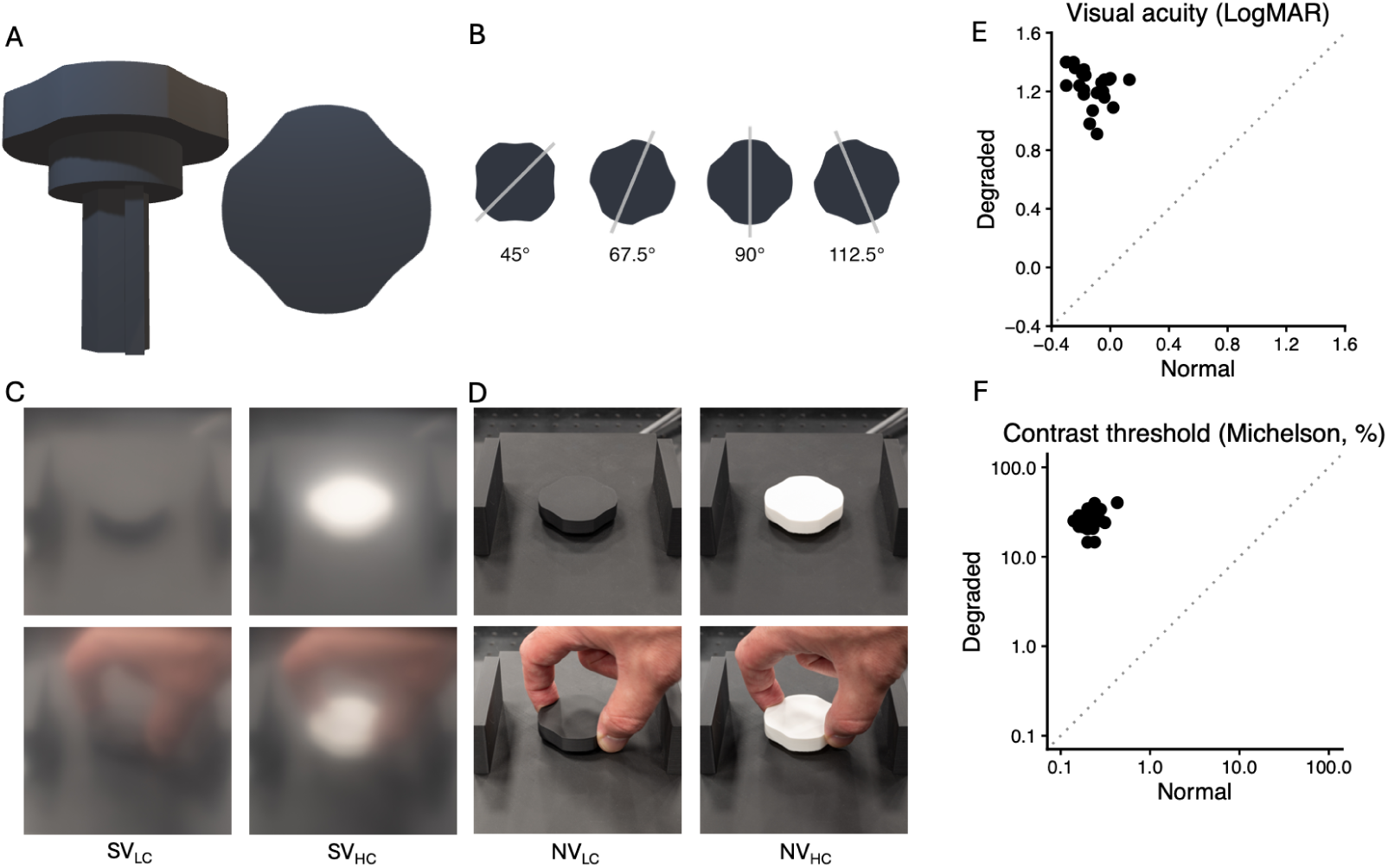
Experimental setup, stimulus configuration, and effects of the low-vision simulation. Participants reached to grasp and lift an object placed in front of them and subsequently returned the hand to the starting position. **A:** Side view of the object (curvilinear octagon with alternating convex and concave arcs) showing the mounting component used to link the object to the motorized rotatory stage (left), and top view of the object (right). The objects were 3D-printed in white and black variants to manipulate stimulus contrast. **B:** Objects were presented at four orientations. **C:** Examples of the visual appearance of the objects under simulated low-vision conditions (SV). The black object on a black background created the low-contrast condition (SV_LC_) whereas the white object on the same background created the high-contrast condition (SV_HC_). **D:** Experimental stimuli and grasping task under normal vision. The black object on a black background created the low-contrast condition (NV_LC_) whereas the white object on the same background created the high-contrast condition (NV_HC_). Finger movements were tracked using markers attached to the thumb and index finger. **E:** Binocular visual acuity (logMAR) measured under normal and degraded viewing conditions using FrACT10 with Sloan letters. **F:** Binocular contrast detection thresholds (Michelson contrast, %) measured using FrACT10 grating stimuli at 2.0 cycles per degree (cpd). Both axes are displayed on a logarithmic (base 10) scale. In panels E and F, each point represents an individual participant, and the dotted line indicates the unity line.

Thumb and index finger movements were recorded at 200 Hz using an OptiTrack motion capture system (NaturalPoint, Inc.). Fingertip positions were tracked by registering each fingertip as the origin of a local coordinate system associated with a rigid body defined by nail-attached passive retroreflective markers (four on the index, five on the thumb), adapted from the MOTOM toolbox.^46^

Stimulus orientation (45°, 67.5°, 90°, and 112.5°, Figure 1B) was controlled using a motorized rotary stage (Velmex Inc.). The target object was a custom 3D-printed curvilinear octagon with alternating convex and concave arcs (top surface: 10 mm thick, maximum diameter: 50 mm, minimum diameter: 45 mm). The object was presented in black or white against a matte-black background to create low- and high-contrast conditions, respectively (Figure 1C–D). The object was designed to afford a stable and easily achievable precision grasping.

Participants completed 10 practice and 160 experimental trials. The experiment employed a 2 × 2 design with Vision (simulated low vs. normal) and Contrast (low vs. high), yielding four conditions (SV_LC_, SV_HC_, NV_LC_, NV_HC_) of 40 trials each, with object orientation randomized within blocks. Simulated low-vision blocks preceded normal-vision blocks to avoid prior visual exposure to the object shape.

Simulated low vision was induced using Cambridge Simulation Glasses, which reduce visual acuity (VA) and contrast sensitivity (CS) through Hampshire Frost diffusing filters,^47^ mounted over habitual spectacles if required. Four filter layers were selected to approximate moderate-to-severe visual impairment^48^ while maintaining object visibility. Occlusion goggles (Red Scientific) were used to prevent vision of the workspace between trials. Binocular VA and contrast detection thresholds were evaluated using the Freiburg Visual Acuity and Contrast Test (FrACT10), with Sloan letters for VA and grating stimuli at a spatial frequency of 2.0 cycles per degree (cpd) for contrast detection thresholds, presented at 80 cm.^49,50^ Contrast thresholds were expressed as Michelson contrast (%), with higher values indicating poorer contrast sensitivity. Mean VA deteriorated from −0.13 ± 0.11 logMAR to 1.23 ± 0.13 logMAR under simulated low vision (Figure 1E). Mean contrast detection thresholds (Michelson contrast, %) increased from 0.27 ± 0.06% to 26.2 ± 6.69% (Figure 1F, both axes on a logarithmic base 10 scale).

### 2.3 Data Analysis

Data were processed in R.^51^ Missing marker samples were interpolated, and position data were smoothed using a Savitzky–Golay filter prior to the calculation of velocity and acceleration. Movement onset and offset were identified using objective kinematic criteria.^12,52^ Movement duration was defined as the time elapsed between movement onset and movement offset. Peak velocity (PV) was defined as the maximum tangential velocity of the thumb during the movement, and time to PV as the elapsed time from movement onset to the occurrence of peak velocity. Path length was calculated as the cumulative three-dimensional distance traveled by the midpoint between the thumb and index finger from movement onset to movement offset. Maximum finger height was defined as the maximum vertical position of the midpoint between the thumb and index finger during the movement. Grasp aperture was defined as the Euclidean distance between the thumb and index finger throughout the movement and was used to determine both maximum grip aperture (MGA), defined as the largest aperture reached during the movement, and final grip aperture (FGA), defined as the aperture at the moment the object was grasped immediately before lift-off. Time to MGA was defined as the elapsed time from movement onset to the occurrence of MGA. Grasp position relative to the object’s center of mass (COM) was quantified as the perpendicular distance between the grasp axis and object center. Grasp axis orientation was defined as the angle of the thumb–index axis in the horizontal plane. Trials with excessive missing data (above 10% of frames), incomplete movements, or extreme final grasp aperture values (above 3 × interquartile range within participant) were excluded. Following pre-processing and outlier removal, 3650 of 3680 trials (99.2%) were retained for analysis.

All dependent variables were analyzed using Bayesian hierarchical models with weakly informative priors (brms package^53,54^). Experimental condition (SV_LC_, SV_HC_, NV_LC_, NV_HC_) was modeled as a fixed effect, with participant-specific random effects. The effects of vision, contrast, and their interaction were quantified using contrasts calculated from the condition-specific posterior distributions. The main effect of vision was defined as the difference between the average of the two normal-vision conditions and the average of the two simulated-low-vision conditions. Similarly, the main effect of contrast was defined as the difference between the average of the two high-contrast conditions and the average of the two low-contrast conditions. The vision-by-contrast interaction was calculated as the difference between the contrast effects under normal vision and simulated low vision. Four Markov chain Monte Carlo (MCMC) chains were run for 4,000 iterations each, including 1,000 warm-up iterations, and posterior distributions were summarized using posterior medians together with 50% and 95% highest density intervals (HDIs). Movement duration, PV, PV time, path length, finger height, MGA, time to MGA, and COM distance were analyzed using hierarchical models estimating condition-specific population means while accounting for between-participant variability. The relationship between MGA and FGA was examined using Bayesian hierarchical regression with condition-specific slopes, allowing the association between these variables to vary across experimental conditions while accounting for participant-specific intercepts and slopes. A complementary Bayesian analysis was performed to characterize grasp contact-point selection. Grasp orientations were transformed into a common reference frame relative to the nearest convex grasp axis, allowing grasps across all object orientations to be pooled. A Bayesian Bernoulli hierarchical model was then used to estimate the probability of selecting a convex rather than a concave grasp region. Finally, for grasps classified as convex or concave, a second Bayesian location– scale model estimated the dispersion of grasp orientations around the center of the selected grasp region separately for convex and concave grasps, thereby quantifying the consistency of grasp placement within each region. Circular kernel density estimates were used to visualize grasp preference distributions and compare the observed data with simulated reference distributions representing random, convex, concave, and equal convex-concave grasp preferences.

## 3 Results

### 3.1 Reach transport

We first examined how visual degradation and contrast influenced the movement towards the object (Figure 2). The temporal structure of the movement remained remarkably stable across conditions (Figure 2A–B). Movement duration was unaffected by simulated low vision or its interaction with contrast. Similarly, peak velocity and timing of peak velocity were largely preserved across conditions (Figure 2C–F). However, under simulated low vision, movements were shorter and lower, with path length reduced by approximately 7.3 mm (95% CrI, 1.4–13.0 mm; Figure 2G and H) and maximum finger height reduced by 5.0 mm (95% CrI, 1.3–8.7 mm; Figure 2I and J) relative to normal vision. Contrast and vision-by-contrast interaction showed no effect. Across all conditions, thumb and index trajectories followed similar global depth-height paths, indicating preservation of the overall transport structure of the movement (Figure 2K). Nevertheless, trajectories under simulated low vision were generally lower, consistent with the reduction in maximum finger height.

**Fig. 2.**
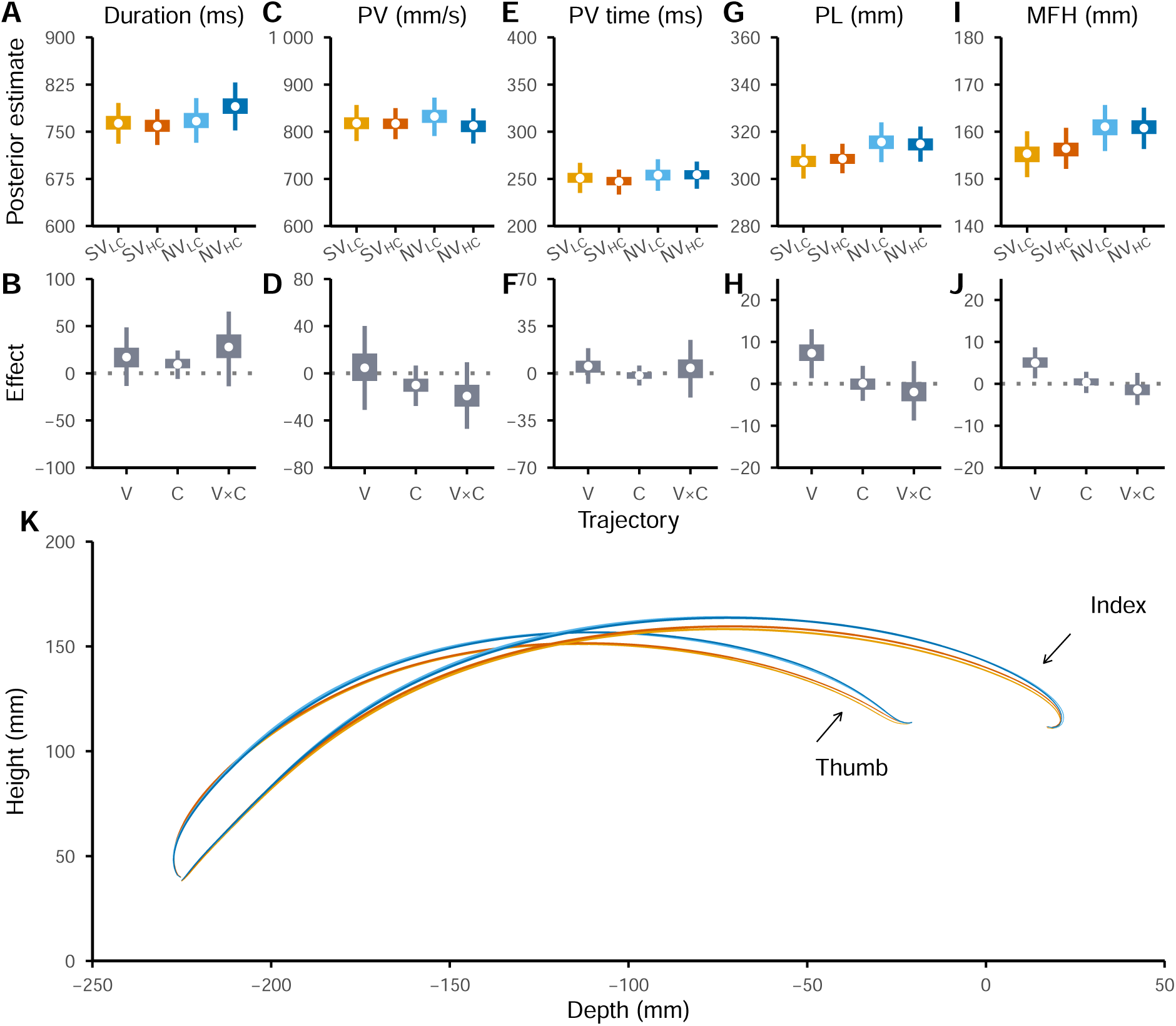
Reach transport. Posterior estimates and corresponding effect estimates derived from Bayesian hierarchical models are shown for movement duration (A, B), peak velocity (PV; C, D), time to peak velocity (PV time; E, F), path length (PL; G, H), and maximum finger height (MFH; I, J). The corresponding effect panels (B, D, F, H, J) show credible differences reflecting the main effects of vision (V), contrast (C), and their interaction (V×C), expressed in the same units as the outcome panels directly above them. White dots indicate posterior medians, boxes indicate the 50% highest density intervals (HDIs), and thin vertical bars indicate the 95% HDIs. Dotted horizontal lines in the effect panels indicate zero difference. Panel K shows the mean movement trajectories of the thumb and index finger across the four experimental conditions: SV_LC_: light orange, SV_HC_: dark orange, NV_LC_: light blue, and NV_HC_: dark blue.

Under normal vision, participants may adopt higher and longer trajectories to reduce object occlusion during the final approach to facilitate online visual feedback. Although these elevated trajectories likely increase movement cost, this may be worthwhile when visual guidance is reliable. In contrast, under degraded vision, participants adopt shorter and lower movements reflecting a more economical transport strategy.

### 3.2 Grasp formation and scaling

To understand how degraded vision affects hand shaping during the approach we analyzed grip aperture evolution over time, as well as MGA magnitude and timing. Across all conditions, grasp aperture followed a similar trajectory, with the hand opening progressively to a peak before closing toward object contact (Figure 3A). MGA remained comparable across vision and contrast conditions (Figure 3B and C), indicating preserved hand shaping and similar safety margins during grasp formation. In contrast, the timing of grip formation was altered by degraded vision. Under simulated low-vision conditions, MGA occurred approximately 27 ms earlier than under normal vision (95% CrI, 1.0–51.7 ms; Figure 3D and E). We found no evidence for contrast or vision-by-contrast interaction effects. This difference may reflect a collision avoidance strategy or result from the higher hand trajectories observed under normal vision, which could delay MGA attainment.

**Fig. 3.**
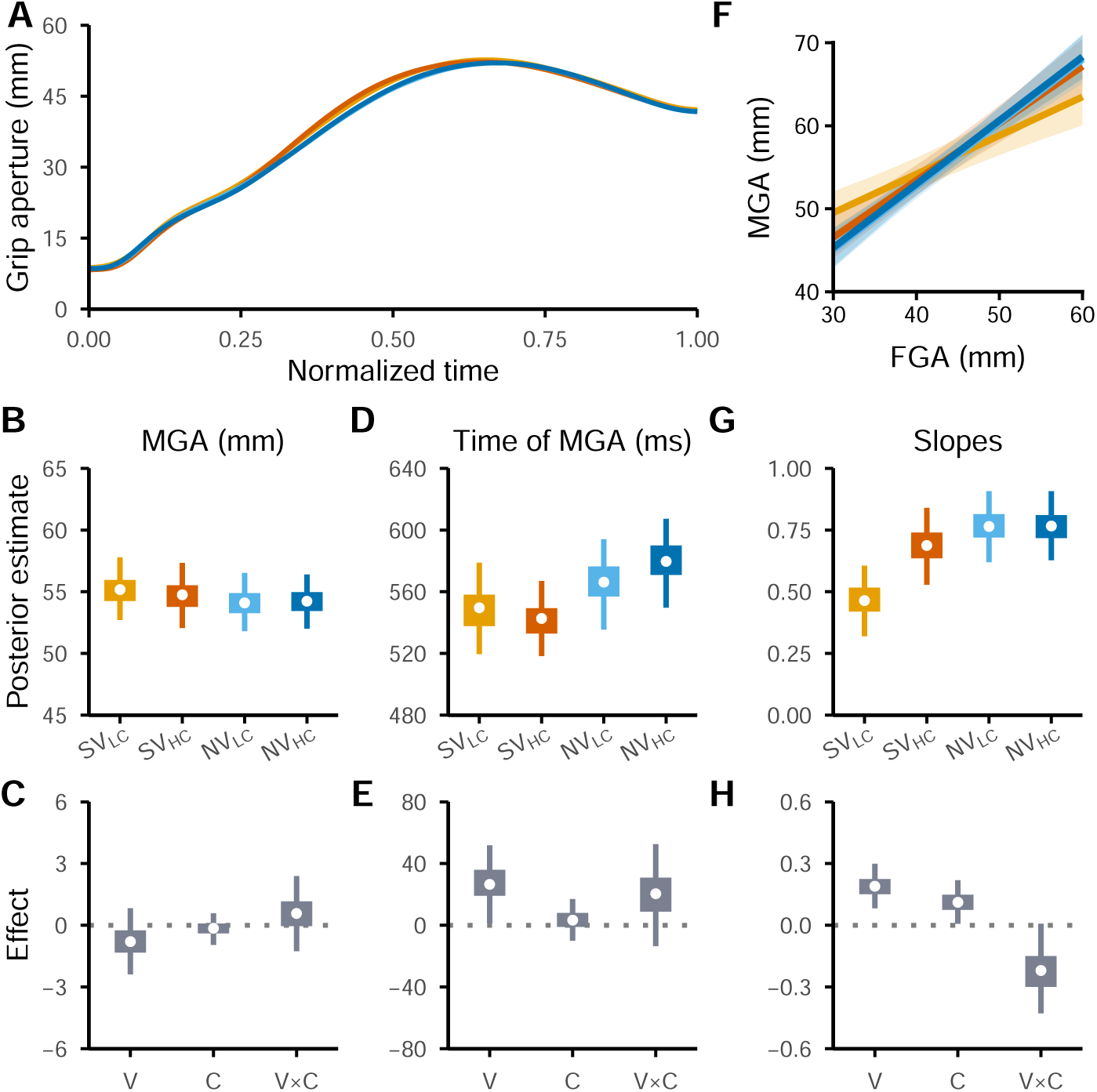
Grasp formation and scaling. Panel A shows mean grip aperture profiles across normalized movement time for four experimental conditions: SV_LC_: light orange, SV_HC_: dark orange, NV_LC_: light blue, and NV_HC_: dark blue. Solid lines represent group mean aperture trajectories. Posterior estimates and corresponding effect estimates derived from Bayesian hierarchical models are shown for maximum grip aperture (MGA, B,C), time of MGA (D, E)). Panels B and D show condition-specific posterior estimates, while panels C and E show the main effects of vision (V), contrast (C), and their interaction (V×C), expressed in the same units as their corresponding outcome panels. Panel F shows fitted population-level predictions of the relationship between MGA and final grip aperture (FGA), with shaded bands representing 95% credible intervals. Panel G shows posterior estimates of the condition-specific MGA-FGA slopes, and panel H shows the corresponding effect estimates for vision, contrast, and their interaction. In panels B–E and G–H, white dots indicate posterior medians, boxes indicate the 50% highest density intervals (HDIs), and thin vertical bars indicate the 95% HDIs. Dotted horizontal lines in panels C, E, and H indicate zero difference

To examine whether grip aperture was appropriately scaled to object geometry under degraded vision, we analyzed the relationship between MGA and final grip aperture (FGA) (Figure 3F). Since FGA depends on the selected contact points, it varies with local object geometry, spanning different diameters. Across all conditions, MGA increased with FGA (Figure 3G), indicating that participants scaled hand opening to object geometry. However, this relationship was weaker under simulated low vision, particularly in the SV_LC_ condition. Vision had a clear impact, supported by secondary Contrast and interaction effects (Figure 3H).

### 3.3 Grasp placement relative to object geometry

To investigate how visual degradation influences grasp spatial organization, we analyzed how the grasp axis aligns with the object’s center of mass and how contact-point selection is affected by local geometry. The distance between the grasp center and the object’s center of mass (COM distance, Figure 4A) increased under simulated low-vision conditions by approximately 0.64 mm (95% CrI, 0.11–1.17 mm; Figure 4B and D), indicating a minor reduction in endpoint accuracy. We found no evidence for contrast or vision-by-contrast interaction effects.

**Fig. 4.**
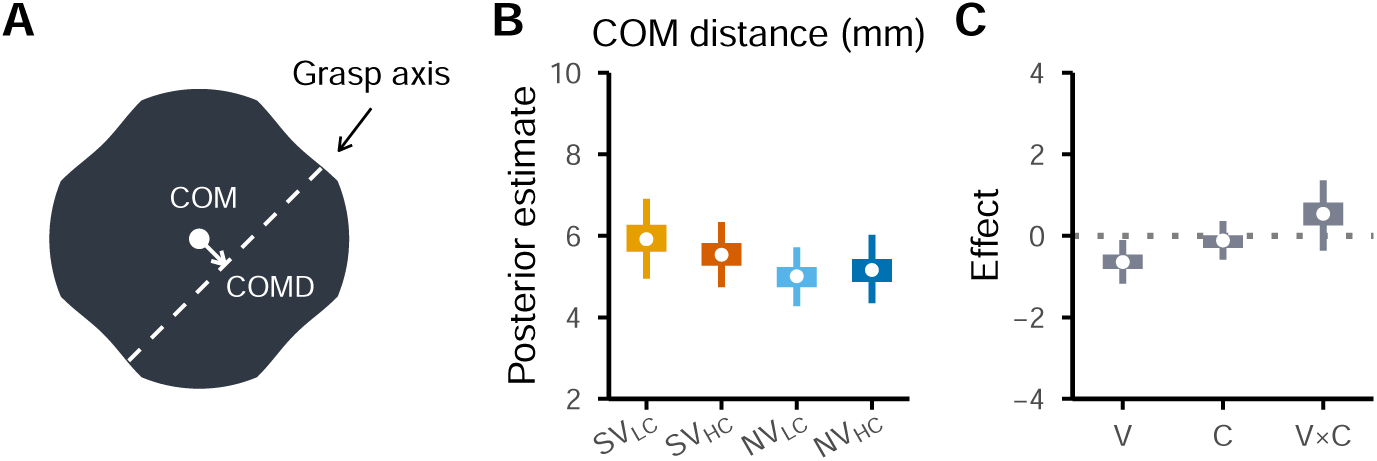
Distance between the grasp axis and the object’s center of mass (COMD). Panel A illustrates COMD, calculated as the perpendicular distance from the object’s center of mass (COM) to the grasp axis defined by the final positions of the thumb and index finger. The dashed white line represents the grasp axis, and the white arrow indicates COMD. Panel B shows the condition-specific posterior estimates of COMD. Panel C shows the main effects of vision (V), contrast (C), and their interaction (V×C), expressed in the same units as panel B. In panels B and C, white dots indicate posterior medians, boxes indicate the 50% highest density intervals (HDIs), and thin vertical bars indicate the 95% HDIs. The dotted horizontal line in panel C indicates zero difference.

To examine geometry-dependent grasp-point selection, we analyzed grasp orientations relative to the object’s geometry. Figure 5A illustrates example grasp axes. Because the object has fourfold rotational symmetry, with convex reference axes repeating every 90°and concave reference axes halfway between them (45°), each measured grasp orientation was expressed as its angular deviation from the nearest convex reference axis (Figure 5B). This transformation maps grasp orientations onto a common angular reference frame. In the polar plots (Figure 5C–J), grasps on convex regions map to the right side (labeled Convex), whereas grasps on concave regions map to the left side (labeled Concave). The closer these grasps are to the center of their respective regions, the closer they align with the right and left directions, respectively. Figure 5C–F show hypothetical reference distributions corresponding to different grasp-selection strategies (random, convex preference, concave preference, and equal preference), while Figure 5G–J show the observed distributions for each experimental condition. Grasp-point selection varied substantially across visual conditions. Under simulated low vision, the distributions approached a more uniform spread (particularly in low contrast). This contrasted sharply with normal vision, where distributions showed clear bimodal peaks corresponding to the centers of the convex and concave grasp regions.

**Fig. 5.**
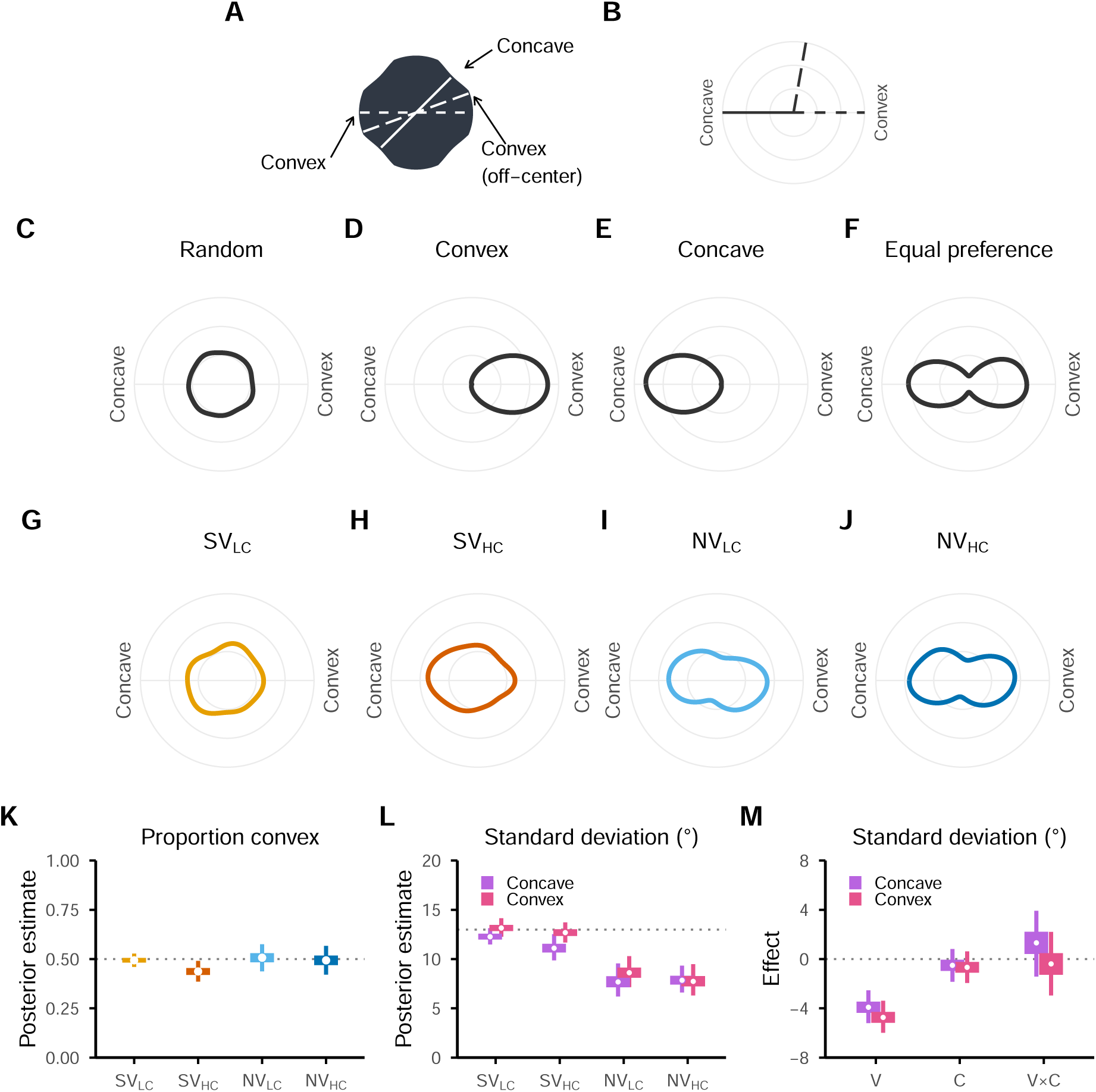
Grasp-point selection relative to object geometry. Panel A illustrates example grasp axes corresponding to convex, concave and off-center convex grasp orientations. Panel B shows the angular representation of the same grasp axes after expressing their orientations relative to the nearest convex reference axis. Because the object has fourfold rotational symmetry, orientations are mapped onto a common range, with grasps on convex regions aligning with the convex reference axis (labeled Convex) and with grasps on concave regions aligning with the concave reference axis (labeled Concave). Panels C–F illustrate hypothetical grasp-orientation distributions: a uniform distribution (C), representing no systematic preference for particular grasp-point selection, equivalent to random grasp-point selection; a single peak centered on convex (D) or concave (E) regions, indicating preferential selection of these regions; and two peaks (F), representing similar preferences for convex and concave regions. Panels G–J show the observed grasp-orientation distributions for simulated low vision–low contrast (SV_LC_, light orange), simulated low vision–high contrast (SV_HC_, dark orange), normal vision–low contrast (NV_LC_, light blue), and normal vision–high contrast (NV_HC_, dark blue), respectively. Panel K shows the model-estimated proportion of convex grasp selections across conditions. Panel L shows condition-specific posterior estimates of grasp-orientation variability (standard deviation, SD) for concave and convex grasp regions. The dotted horizontal line represents the SD expected under a uniform distribution of grasp orientations. Panel M shows the estimated effects of vision (V), contrast (C), and their interaction (V×C) on grasp-orientation variability. In panels K–M, white dots indicate posterior medians, boxes indicate the 50% highest density intervals (HDIs), and thin vertical bars indicate the 95% HDIs. The dotted horizontal line in panel M indicates zero difference.

Although the overall proportion of convex selections remained close to 0.5 across conditions, indicating no systematic preference for either convex or concave regions (Figure 5K), the SV_HC_ condition showed a modest shift toward concave selections. The consistency of geometry-dependent grasp-point selection was estimated using the variability of grasp orientations within the convex and concave object regions (Figure 5L–M). Greater variability indicates that grasp locations were more widely dispersed, reflecting less consistent geometry-dependent selection. Variability in simulated low vision increased by 3.91° (95% CrI, 2.55–5.20°) for concave and by 4.74° (95% CrI, 3.45–6.04°) for convex grasp locations relative to normal vision. The effects of contrast were comparatively small.

These findings indicate that simulated low vision eliminated the influence of object geometry on grasp-point selection, resulting in grasp orientations that approached a random distribution, with a modest attraction toward the concave regions in the SV_HC_ condition. In contrast, when object geometry was visually accessible under normal vision, grasp axes clustered distinctly at the centers of both convex and concave regions, demonstrating no systematic preference for either.

## 4 Discussion

The present study examined how degraded visual input influences grasp movements and object-geometry dependent grasp-point selection. While the overall structure of the action was preserved, the extent to which movements were tuned to the spatial properties of the object was strongly impacted under simulated low-vision conditions.

### 4.1 Reach transport

Movement duration, peak velocity, and the timing of peak velocity were largely preserved under degraded visual conditions. Thus, the temporal structure of the reach remained stable, indicating that acute visual degradation did not affect movement execution.

This contrasts with studies of clinical visual impairment, which have often reported broader changes in reach-to-grasp kinematics, including longer movement durations, prolonged deceleration phases, reduced peak velocities, and increased online corrections.^20,23,26–28,55^ Thus the impact of reduced vision depends on the nature of the visual deficit. In chronic visual impairment, movement control may be shaped by spatially localized visual field loss, reduced reliability of visual feedback, and long-term adaptation. For example, central scotomas may obscure both the object and the moving hand during the reach, increasing uncertainty about object location and hand position and placing greater demands on online corrections.

In contrast, the present study used an acute and global reduction of visual quality. Although this manipulation substantially reduced VA and CS, it did not create a localized visual field defect. Participants under simulated low vision showed shorter and lower trajectories, while the temporal structure of the reach remained largely preserved. These findings suggest that when visual information is degraded but still available, participants rely on a stable motor plan while adopting a more conservative movement strategy. This interpretation is consistent with evidence from simulated visual degradation and long-term stereoblindness showing that visuomotor control adapts according to the reliability, duration, and spatial characteristics of available visual information.^30,56–58^ It is also consistent with evidence that visual information contributes to the spatial calibration of reaching movements and that degraded visual input can promote more constrained movement strategies under conditions of uncertainty.^20,27,59–62^

### 4.2 Grasp formation and scaling

Grasp formation showed a similar dissociation between preserved structure and altered temporal organization. MGA remained largely unchanged across conditions, indicating that participants maintained appropriate safety margins when shaping the hand to grasp the object. In contrast, MGA occurred earlier under low vision, suggesting a shift in the coordination between transport and grasp components.

This pattern is broadly consistent with previous studies showing that MGA is often preserved despite substantial impairments in movement timing and execution in visually impaired populations.^20,26^ At the same time, degraded visual input has been associated with increased reliance on non-visual feedback during the later stages of grasping and reduced precision in hand–object interactions.^23,28,55^ The earlier occurrence of MGA observed here is consistent with a reduced contribution of late-stage visual feedback, suggesting greater reliance on feedforward control of hand shaping.

The effects of degraded vision became more apparent when considering how grip aperture was scaled to object geometry. Although MGA scaled systematically with FGA in all conditions, this relationship was weaker under simulated low vision, indicating reduced sensitivity to object geometry. Thus, hand shaping remained tuned to object structure, but this tuning became less precise when visual information was degraded.

### 4.3 Grasp placement relative to object geometry

Simulated low vision increased the distance between the grasp axis and the object’s center of mass, indicating reduced grasp-centering accuracy. Because grasp centering maximizes object stability and minimizes torque,^33,63^ participants likely relied more on intrinsic motor tendencies than object-specific information. Similar shifts toward default contact-point distributions have been reported when object properties cannot be estimated reliably.^36^ Such reductions in spatial accuracy may contribute to the difficulties experienced by individuals with low vision during everyday object manipulation, particularly when successful interaction depends on accurate finger placement relative to object features.

Visual degradation most prominently affected geometry-dependent contact-point selection. Participants showed no clear preference for convex over concave regions under either normal or degraded vision. However, visual conditions markedly affected where within these regions contact points were selected. Under normal vision, grasp points were consistently concentrated at the centers of convex and concave regions. Under degraded vision, grasps distributed more uniformly around the object’s perimeter, indicating that object geometry no longer systematically guided selection. This loss of spatial precision may contribute to the difficulties experienced by individuals with visual impairment during tasks requiring precise finger placement and highlights functional consequences that conventional measures of visual performance may not capture.

These findings should be interpreted with a few important caveats in mind. While simulated low vision cannot fully replicate heterogeneous visual deficits or long-term adaptations associated with clinical visual impairment, it offers the significant advantage of providing precisely controlled visual degradation. Additionally, future studies will need to generalize these findings to everyday object manipulation beyond a single object geometry and a controlled grasping task.

### 4.4 Conclusions

Simulated low vision weakened the link between object geometry and precise finger placement while preserving the overall organization of grasping. Successful object manipulation can therefore mask a loss of precision in visually guided action. These findings highlight the potential of grasping movement measures to complement conventional clinical assessments providing a finer understanding of how visual impairment impacts everyday interactions with objects.

## Acknowledgments

This work was partially supported by the NYUAD Center for Artificial Intelligence and Robotics, funded by Tamkeen under the NYUAD Research Institute Award CG010, and, by the NYUAD Center for Brain and Health, funded by Tamkeen under the NYUAD Research Institute Award CG012.

This work was presented as an oral presentation entitled “Contact-point selection in grasping under degraded vision” at the 2026 Vision Sciences Society (VSS) Annual Meeting, held May 15–19, 2026, in St. Pete Beach, Florida, USA.

Disclosure: **C. Lanca**, Eyerising (C), Essilor Portugal (C); **Z. Derzsi**, none; **R. Volcic**, none.

## Data availability

Data are available at https://doi.org/10.5281/zenodo.24268854.

## Credit author statement

**Carla Lanca:** Designed the experiment, Conducted the experiment, Analyzed/interpreted data, Wrote the article. **Zoltan Derzsi:** Designed the experiment; Proofed/revised article. **Robert Volcic:** Designed the experiment, Provided materials, Analyzed/interpreted data, Proofed/revised article.

